# Exploring the impact of variability in cell segmentation and tracking approaches

**DOI:** 10.1101/2023.11.24.568598

**Authors:** Laura Wiggins, Peter J. O’Toole, William J. Brackenbury, Julie Wilson

## Abstract

Segmentation and tracking are essential preliminary steps in the analysis of almost all live cell imaging applications. Although the number of open-source software systems that facilitate automated segmentation and tracking continue to evolve, many researchers continue to opt for manual alternatives for samples that are not easily auto-segmented, tracing cell boundaries by hand and re-identifying cells on consecutive frames by eye. Such methods are subject to inter-user variability, introducing idiosyncrasies into the results of downstream analysis that are a result of subjectivity and individual expertise. Such methods are also susceptible to intra-user variability, meaning findings are challenging to reproduce. Here we demonstrate and quantify the degree of intra- and inter-user variability in manual cell segmentation and tracking by comparing the phenotypic metrics extracted from cells segmented and tracked by different members of our research team. Furthermore, we compare the segmentation results for a ptychographic cell image obtained using different automated software and demonstrate the high dependence of performance on their imaging modality optimisation. Our results show that choice of segmentation and tracking methods should be considered carefully in order to enhance the quality and reproducibility of results.

## Introduction

With microscopic imaging becoming an essential step in almost all areas of biological research, there is an increased demand for image analysis tools that provide reliable phenotype quantification.^1^ A particular challenge in the face of image analysis is *segmentation*, a means of distinguishing objects from background. Objects of interest vary depending on application, with examples including single-cells within 2D microscopy images,^2^ tumours within magnetic resonance images^3^ or tissue from histopathological whole-slide images.^4^ At present, many studies make use of manual segmentation to identify regions of interest (ROIs), though recent research has highlighted the issue of both inter- and intra-operator variability in manual segmentation and the impact this has upon reproducibility and biological conclusions.^5, 6, 7, 8^ The subjectivity of manual segmentation and the time required to identify ROIs by hand has motivated a shift to automated approaches that are standardised, delineated and high-throughput.^9^ This has had a particular impact in clinical settings where automated approaches are now used to make diagnoses through computer-aided diagnosis (CAD) systems.^10^

Although providing benefits in terms of reproducibility, automated segmentation approaches still involve some degree of user interactivity, with manual segmentation often considered the gold standard against which to benchmark the results of automated segmentation.^11, 12^ Furthermore, model-based segmentation approaches, such as U-Nets, make use of manually segmented ROIs for ground truth training sets.^13^ Automated approaches are also often specific to certain imaging modalities, with models learning characteristics of such images and using this knowledge to accurately identify ROIs whilst excluding background and image artefacts. For all of these reasons, choice of segmentation approach prior to quantification from microscopy images is a hugely important step that should not be overlooked and should instead be carefully considered.

There exist several established, open-source software packages for the task of segmenting cells from 2D microscopy images, each with their own strengths and limitations.^14^ Studies have been carried out to benchmark such software against one another based on accuracy of segmentation and ease of usability, though notably fluorescence images that have been used to trial the majority of these software. With the emergence of label-free imaging, efforts are now being made to form ground truth data sets of label-free cells to assist in the training of segmentation models that are specific to label-free imaging techniques such as phase-contrast, bright-field and quantitative phase imaging (QPI).^15^ More recent cell segmentation software such as CellPose^16^ emphasise their focus on developing models that are adaptable to new imaging techniques.

Here we demonstrate the direct impact that manual segmentation can have on downstream analysis by quantifying the inter- and intra-user variability of phenotypic metrics extracted from manually obtained ROIs. Moreover, we show that manual cell tracking also results in large variability in the extraction of motility metrics due to subjectivity in the choice of cell centroid position as well as re-identification of cells on consecutive frames. By comparing the results of automated and manual cell segmentation and tracking, we demonstrate greater reproducibility for a panel of automated packages, but large inter-software variability arising from differences in imaging modality optimisation.

## Methods

### Cell lines

The MDA-MB-231 breast cancer cell line used within this study was a gift from M. Djamgoz, Imperial College, London. The molecular identity of this cell line was verified by short tandem repeat analysis.^17^ Thawed cells were sub-cultured 1-2 times prior to discarding and thawing a new stock to ensure that the molecular identity of cells was retained throughout. Cells were cultured in Dulbecco’s modified eagle medium supplemented with 5% fetal bovine serum (FBS) and 4mM L-glutamine.^18^ FBS was filtered using a 0.22*µ*m syringe filter prior to use to reduce artefacts when imaging. Cells were incubated at 37*^◦^*C in plastic filter-cap T-25 flasks and were split at a 1:6 ratio when passaged. No antibiotics were added to cell culture medium.

### Image acquisition

Cells were placed onto the Phasefocus Livecyte 2 (Phasefocus Limited, Sheffield, UK) to incubate for 30 minutes prior to image acquisition to allow for temperature equilibration. One 500*µ*m x 500*µ*m field of view per well was imaged to capture as many cells, and therefore data observations, as possible.

### Manual segmentation and tracking

In order to compare inter-user variability, members of our research group with different research expertise performed the same segmentation and tracking tasks. Manual segmentation was performed using ImageJ’s freehand area selection tool to trace around individual cells and selections were added to ImageJ’s ROI manager using Edit > Selection > “Add to Manager”. Morphological features were then extracted from each ROI using ImageJ’s Measure function, where all features were selected from the available list provided in Analyze > Set Measurements. A full list of extracted features together with definitions is provided within the ImageJ documentation.

ImageJ’s MTrackJ plugin was used for manual tracking of cells. Each cell was tracked for 50 frames by clicking the cells’ position on consecutive frames, with X and Y coordinates from all final trajectories exported for feature extraction. R’s **trajr**^19^ package was used for the extraction of several features to characterise cell trajectories. The list of extracted metrics comprises: mean and standard deviation of cell speed, sinuosity, trajectory straightness index, mean and standard deviation of track length, trajectory distance and mean trajectory turning angle.

Throughout the paper, we refer to the results obtained by different research team members by a number so that the effect of differences in research experience can be considered.

### Automated segmentation and tracking

#### FIJI segmentation

Fiji^20^ is an open-source software for processing of biological microscopy images. It offers accessible implementation of a range of traditional image processing algorithms, with the user setting subjective thresholds throughout the course of a segmentation workflow. Initially a rolling ball algorithm with radius size 30px was applied to the image for the removal of background noise and imaging artefacts. The image was then smoothed using Gaussian blurring with a standard deviation of 3 to further denoise the image. The image was then converted to a binary mask where each pixel was classified as either background (black within the binary mask) or cell (white within the binary mask) and the watershed method used to separate connected cells ahead of final segmentation by using the “create selection” feature in Fiji for cell boundary detection. A number of alternative workflows were assessed in Fiji such as the use of “find edges” and thresholding prior to binary mask conversion but these methods proved unsuccessful, the details included here describe the workflow identified as being most successful for the segmentation of our images.

#### Livecyte segmentation

Phase Focus’ CATbox software facilitates automated cell segmentation by allowing the user to create a unique image processing recipe for each experiment. This recipe is produced through user-imposed thresholds on a range of image processing techniques allowing for live inspection of segmentation results. A rolling ball algorithm was applied to the image with radius 29.24 *µ*m and a height of 2, to remove background noise and imaging artefacts. The image was then smoothed with a parameter of 19.11*µ*m ahead of seedpoint detection. Local pixel intensity maxima identified from the smoothed image were used as seedpoints with an intensity threshold of 0.8. Nearby seedpoints were then consolidated by combining two seedpoints if there was a path between them that did not change in phase by more than a threshold of 1.09. Thresholds were then applied to distinguish between the pixel intensities of cell and background. A background threshold of 0.05 was used, any pixel with intensity below this threshold was classified as background. A feature threshold (for detection of cells) of 0.6 was used, any pixel above this threshold was classified as belonging to a cell. Pixels that did not meet these requirements were then scaled to a range of 0-1 and formed a “grey area” where pixel classification was uncertain. A biased fuzzy distance transform^21^ was then used to classify “grey area” pixels with a fuzzy distance threshold of 0.2 *µ*m and a phase weighting of 6. Final segmentation results were then exported as .roi files. All thresholds were selected subjectively by assessing live segmentation and revising thresholds to yield optimum results.

#### Icy segmentation

Icy^22^ is a free, open-source image processing software. A key component of Icy is its public repository for the sharing of analysis plug-ins and workflows. The “HK-means” plug-in was utilised for initial segmentation of the cells. This plug-in performs *K*-means clustering of the image pixel intensity histogram to distinguish between background and cells, and then uses size filtering to split undersegmented cells. A Gaussian pre-filter was applied with standard deviation set equal to 5 for image smoothing prior to cell edge detection. *K* was set to 2 to establish two classes of pixel intensity: background and cell. Area size filters were then imposed with a minimum threshold of 100px and a maximum of 5000px, detected objects with areas outside of this range were discarded. The “active contours” plug-in was then used with default parameter settings for closer sculpting of ROIs to cell edges. Finally the “split ROI” plug-in was used to separate cells that were originally undersegmented. This plug-in is semi-automatic and relies on the user to enter how many cells are incorrectly clustered within a group before selecting the group that needs separating. This method was applied to all instances of undersegmentation but some cells were unable to be corrected and were therefore left as is in the final segmentation.

#### CellProfiler segmentation

CellProfiler^23^ is an open-source software for the analysis and measurement of cell images in which pipelines can be constructed by the user for tailored cell segmentation and tracking. Cells were detected using the “identify primary objects” module with a minimum expected object diameter of 10px and a maximum of 50px. Objects detected outside of this range and objects that were touching the borders were discarded. The default image processing settings associated with this module were used, hence the image was globally thresholded prior to segmentation. A minimum cross entropy thresholding method^24^ was used to minimise the average error in pixel classification and Gaussian smoothing with a standard deviation of 1 was performed prior to final segmentation. CellProfiler allows the user to select which method should be used to distinguish and separate clumped objects, in this case intensity was selected as opposed to the use of shape. After running the completed pipeline, segmentation masks exported as .tiff files. To convert image masks into ImageJ ROIs, the masks were read into ImageJ and an ROI of the full mask created using Edit > Selection > Create Selection. This selection was then added to the ROI manager where the “Split” function was used to create individual ROIs from the full mask selection.

#### CellPose segmentation

CellPose^25^ is an open-source, deep learning-based segmentation tool together with pre-trained models for the segmentation of cells, nuclei and tissue sections without the need for parameter fine-tuning. The underlying algorithm involves the transformation of ground-truth cell segmentation into horizontal and vertical flow representations that can be predicted by a neural network to restore accurate segmentation of even unusual cell shapes. CellPose was utilised for cell segmentation using FIJI’s CellPose-TrackMate plugin. The pre-trained cytoplasm model was used for detection of cell boundaries with expected cell diameter of 10 microns.

#### Statistical analyses

All tests of statistical significance within this study were performed using Graphpad Prism 9.1.0 (GraphPad Software, San Diego, CA). Data were tested for normality using the D’Agostino & Pearson test. Parametric tests (t-tests and F-tests) were used where suitable with non-parametric Mann-Whitney U-tests in place of t-tests where data did not follow a normal distribution. For comparison of three or more populations, ANOVA was used when data followed a normal distribution or Kruskal-Wallis tests if not. Tukey’s post-hoc test and Dunn’s multiple comparisons were performed following ANOVA and Kruskal-Wallis respectively. Results were considered significant if *p <* 0.05. Levels of significance used: * *<* 0.05, ** *<* 0.01, *** *<* 0.001, **** *<* 0.0001. Full details of statistical tests used for each analysis are provided in the figure legend for the corresponding figure. Sørensen–Dice coefficients to quantify the similarity of segmented regions were computed using MATLAB’s dice() function.

## Results

### Inter-user variability in manual cell segmentation

Within the current study, researchers 1, 2 and 5 are researchers who have received in-depth microscopy training, including the acquisition and interpretation of microscopy images. In comparison, researchers 3 and 4 are not experienced microscopists but have extensive experience working with the imaged cell line, MDAMB-231. All 5 researchers were set the task of manually segmenting three specified cells from a single microscopy image of MDA-MB-231 cells, the three cells are highlighted in **Figure 1(a)**. As the appearance of the image could differ with the researchers’ computer monitor and display preferences, we provided a contrast enhanced cell image in which cell protrusions not clearly visible within the original image could be observed (**Figure 1(a)**). The extent of protrusions that could be observed by researchers would likely be impacted by their monitor and display preferences.

**Figure 1:**
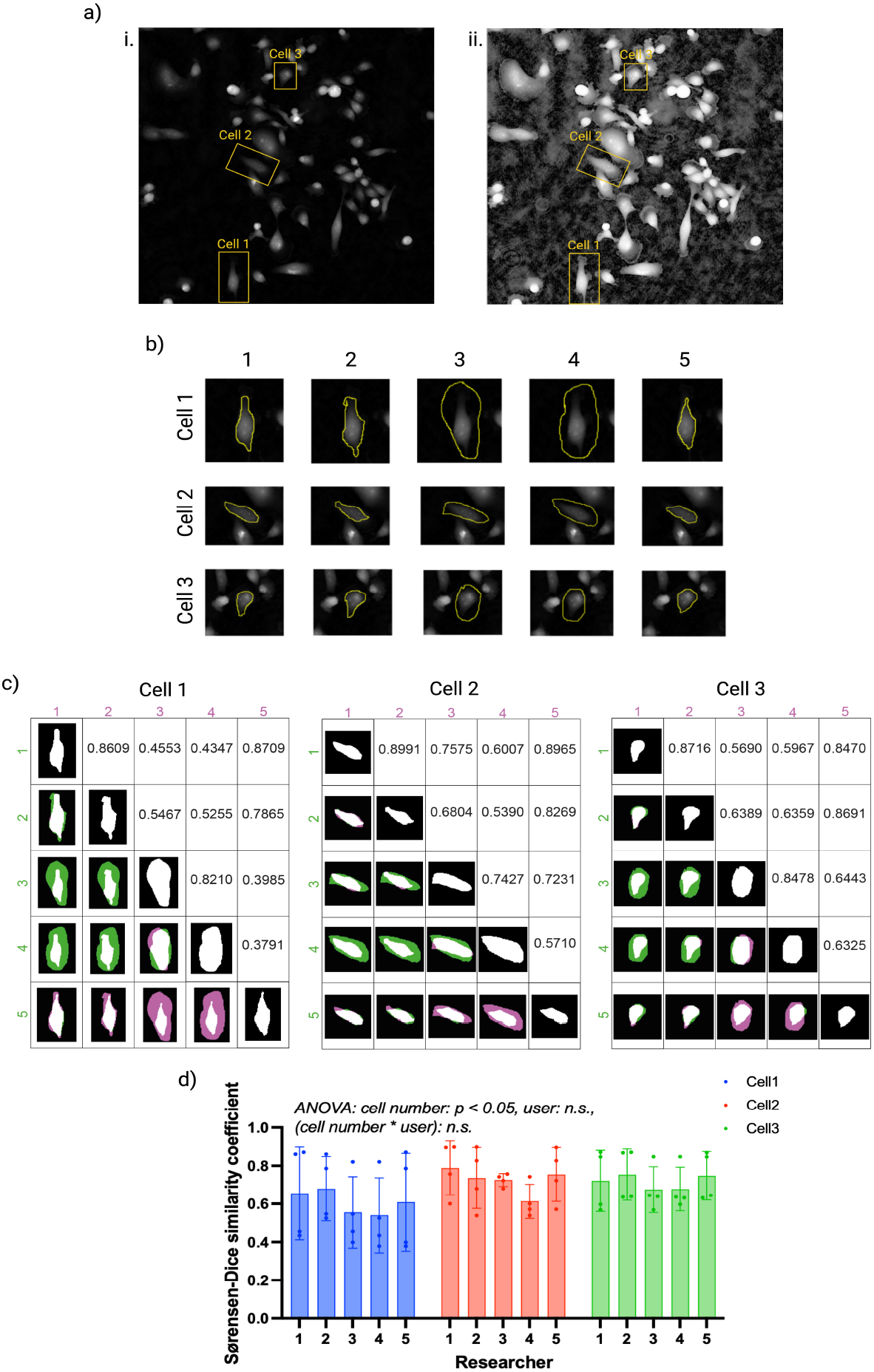
Inter-user variability in manual cell segmentation. **(a)** i. Phase image of MDA-MB-231 cells with the three cells to be manually segmented highlighted and labelled. ii. The cell image with contrast enhanced to display cell protrusions that are not clearly visible within the original cell image. The three cells to segment are highlighted and labelled. **(b)** Manual segmentation of each cell performed by each researcher. Inter-user variability can be observed from differences in outlined ROIs for each cell. **(c)** Sørensen–Dice coefficients to quantify the degree of similarity of pair-wise manual segmentation. Images show segmentation maps where black represents background, white represents segmentation overlap and green and pink represent differences in pair-wise segmentation. Areas in green appear in the segmentation map from the researcher listed in the row position, but not the researcher listed in the column position and viceversa for areas in pink. **(d)** Sørensen–Dice coefficients coefficients, grouped by researcher and coloured by cell number. Bars are representative of group means and error bars are representative of standard deviations to display the spread of coefficients. A two-way ANOVA was performed, obtaining *p* values of 0.0498, 0.3558 and 0.9944 for cell number, researcher, and their interaction, respectively.

Inter-user variability in manual segmentation was observed from each researcher’s identified ROIs (**Figure 1(b)**). Researchers 1, 2 and 5 tended to closely sculpt the brightest intensity pixels, treating the cell boundary as the region around the cell in which bright intensity pixels are neighboured with dark intensity background pixels. Researchers 3 and 4, on the other hand, tended to include a large number of dark intensity pixels within the areas they determined to be the cell interior, despite the resulting ROIs differing in morphology to that expected of an MDA-MB-231 cell. It is possible that researchers 3 and 5 were considering the brightest intensity pixels as cell nuclei, and darker intensity pixels surrounding these areas as cytoplasm. This could be due to their experience with nucleic stains such as DAPI rather than label-free imaging in which whole cells can be visualised.

Sørensen–Dice coefficients confirmed the presence of inter-user variability in segmentation across all three cells, with higher coefficients indicative of significant overlap in segmentation results and lower coefficients representing manual segmentation that differs (**Figure 1(c)**). The Dice coefficients show the impact of research expertise on manual cell segmentation results, with researchers with similar expertise (researchers 1, 2 and 5 or researchers 3 and 4) obtaining high pair-wise coefficients in comparison to the coefficients obtained for researchers with differing expertise.

For each cell, bar charts showing the distribution of Dice coefficients for each researcher’s segmentation data are plotted in **Figure 1(d)** to aid visualisation. Two-way ANOVA revealed a statistically significant relationship between Dice coefficients and cell number, indicating greater inter-user variability in manual segmentation for certain cells. In comparison, no relationship was found between Dice coefficients and researcher number, suggesting that inter-user variability affected all researchers as opposed to one particular researcher. No statistical significance was found between Dice coefficients and the interaction between cell number and researcher, suggesting that inter-user variability is not related to particular cells.

### Intra-user variability in cell segmentation

The segmentation of the three cells was repeated 5 times and the intra-user variability of each researcher’s 5 attempts is shown in **Figure 2(a)**. Visual inspection of each researchers’ segmentation attempts showed that some were more consistent in their segmentation approach, repeatedly outlining similar ROIs. This was the case for researchers 1, 2 and 5 who all aimed to closely sculpt the bright intensity pixels that neighbour dark intensity background pixels. In comparison, researchers 3 and 4 tended to show greater variance in their segmentation attempts where ROI selection, and therefore expected cell shape, was not guided as much by the bright intensity pixels.

**Figure 2:**
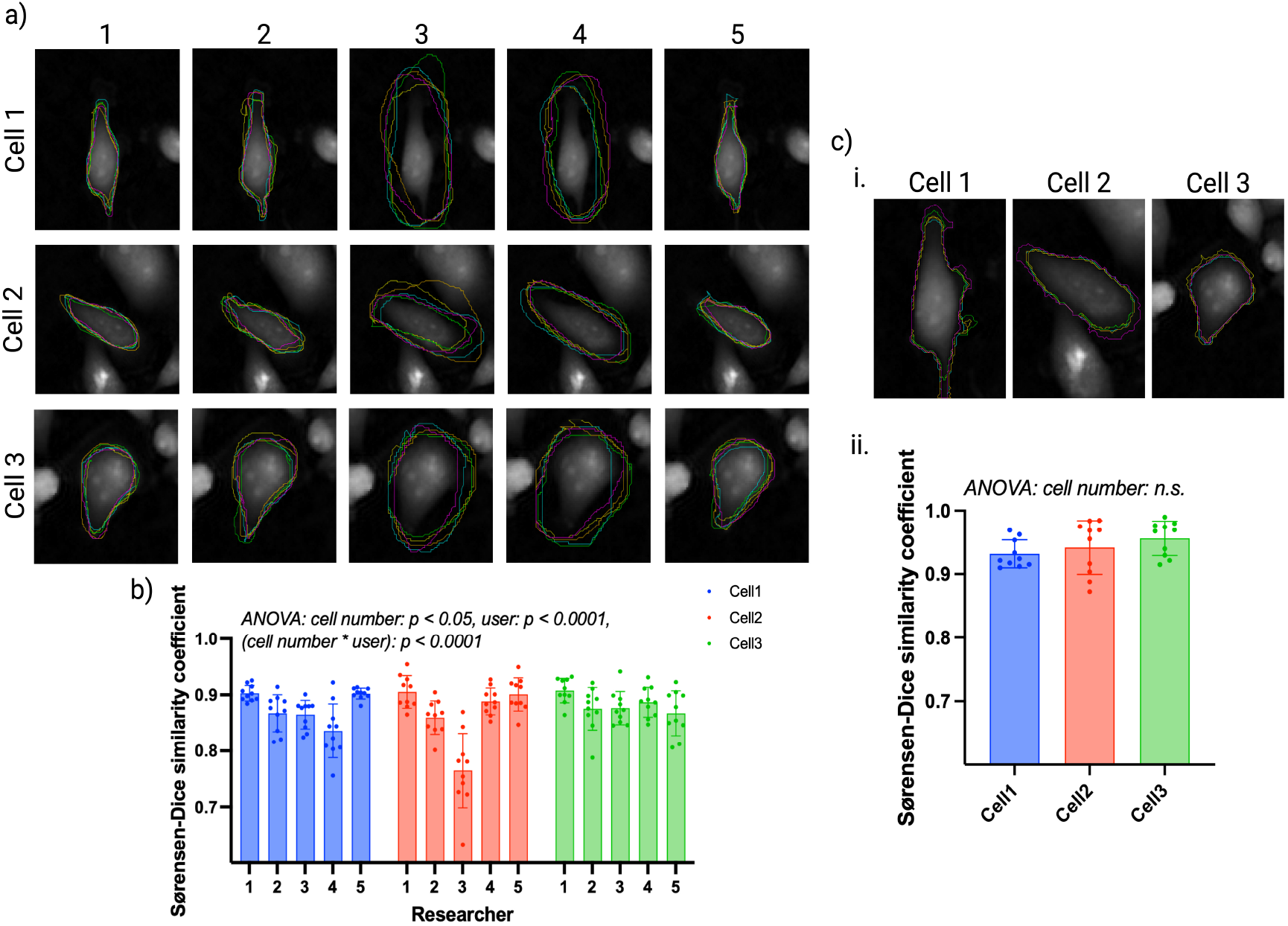
Intra-user variability in manual cell segmentation. **(a)** Manual segmentation of each cell performed by each researcher, each colour represents one of the five segmentation attempts performed. Intra-user variability can be observed from differences in outlined ROIs. **(b)** Sørensen–Dice coefficients, grouped by researcher and coloured by cell number. Bars show group means with error bars representing standard deviations. Two-way ANOVA showed that cell number, researcher and their interaction all have a statistically significant relationship with intra-user Dice coefficients, contributing to 3%, 25% and 28% of the variance prevalent within the data set, respectively. **(c)** i. Automated segmentation of each cell performed using Phasefocus CATbox. Each colour represents one of the five segmentation results obtained from the same cell image processed independently. ii. Sørensen–Dice coefficients, coloured by cell number. ANOVA showed no statistically significant relationship between Dice coefficients and cell number, indicating consistency in segmentation results obtained from Phasefocus CATbox irrespective of cell number.

Intra-user Dice coefficients were calculated and the results for each researcher and each cell are displayed in **Figure 2(b)**. Two-way ANOVA showed a highly significant relationship between intra-user Dice coefficients and researcher number, confirming greater consistency in manual segmentation for some researchers. Differences between researchers accounted for 25% of the total variance present within the data set. A statistically significant relationship was identified between intra-user Dice coefficients and cell number, indicating that intra-user variability was more prevalent for certain cells in comparison to others, though cell number only accounted for 3% of the total variance within the data set. High statistical significance was also found between intra-user Dice coefficients and the interaction between cell number and researchers, showing that maintaining consistency in segmentation proved more challenging with some cells than others for certain researchers. The interaction between cell number and researcher accounted for 28% of the total variance present within the data set.

To assess whether automated approaches to cell segmentation would provide less variable, and therefore more reproducible, results, the same three cells were segmented using the Phasefocus CATbox system. To replicate the potential variance introduced by researchers, the cell image was independently processed five times by an expert with extensive experience in working with ptychographic images and the MDA-MB-231 cell line, and ROIs of the three cells were obtained from each of the five processed images. Visual inspection of these ROIs showed greater consistency in comparison to the intra-user results of the researchers, across all three cells (**Figure 2(c)i**), with differing image processing pipelines resulting in subtle differences in segmentation. Dice coefficients were calculated to assess the similarity of ROIs obtained from the five processed images, the results are displayed in **Figure 2(c)ii**. One-way ANOVA revealed no statistically significant difference in Dice coefficients as a result of cell number (*p* value = 0.2407), indicating greater consistency in segmentation results irrespective of cell number for Phasefocus segmentation.

### The impact of cell segmentation on extracted metrics

As the morphological metrics that can be extracted from cell images rely on initial segmentation of individual cells, accuracy and consistency in identification of ROIs is vital for meaningful downstream analyses. We hypothesised that the inter- and intra-user variability present in manual segmentation would also be prevalent in the morphological metrics extracted from the ROIs. We therefore extracted several morphological metrics describing cell shape, size and texture from the ROIs in **Figure 2(a)**, as well as the Phasefocus CATbox segmentation results (described from now on as Livecyte) in **Figure 2(c)**, and assessed their variability, both inter- and intra-user. Cell area, circularity and mean gray value were chosen as representative descriptors of cell size, shape and texture respectively and their varying values can be seen in **Figure 3(a)**. ANOVA of all three metrics identified statistically significant differences in size, shape and texture of cells as a result of inter-user manual segmentation with *p* values < 0.0001 for all three metrics.

**Figure 3:**
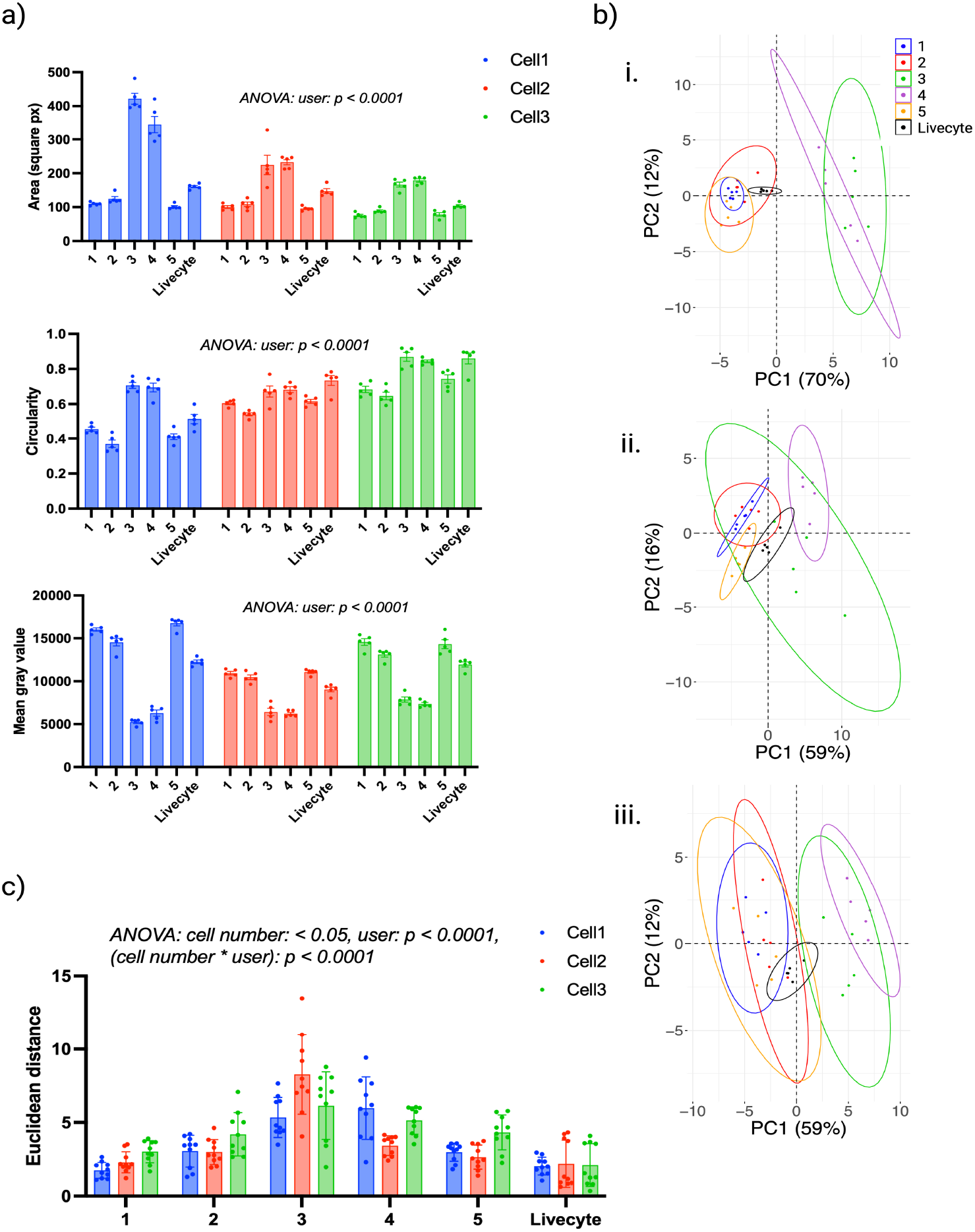
The impact of cell segmentation on extracted metrics. **(a)** Distributions for cell area (top), circularity (middle) and mean gray value (bottom) extracted from different manual segmentation results and Livecyte automated segmentation. Plots are grouped by researcher and coloured by cell number. Bars show group means with error bars representing SEM. ANOVA results show high statistical significance across all three metrics, demonstrating the impact of researcher-specific segmentation on extracted morphological metrics. **(b)** PCA scores plots for i. cell 1, ii. cell 2 and iii. cell 3 obtained from morphological metrics extracted from each manual segmentation and Livecyte automated segmentation. Points are coloured according to researcher and 95% concentration ellipses are displayed. Small and large ellipses are representative of low and high intra-user variability in extracted metrics respectively. **(c)** Pair-wise Euclidean distances between extracted data vectors, grouped by researcher and coloured by cell number. ANOVA revealed statistical significance differences in Euclidean distances as a result of cell number, researcher and the interaction between the two. Bars show group means and error bars represent standard deviations to display the spread of coefficients.

Post-hoc analysis was performed using Tukey’s multiple comparisons test and the full results for cell area, circularity and mean gray value are provided in **Supplementary Table 1, 2** and **3** respectively. Notably, researchers 1, 2 and 5 frequently displayed no significant differences in extracted metrics when compared with one another (with the exception of researchers 2 and 5 scoring a *p* value of 0.0356 for circularity), but did consistently differ significantly from researchers 3 and 4 across all metrics. Furthermore, researchers 3 and 4 frequently displayed no significant differences between one another. Differences in research expertise between researchers 1, 2 and 5, and researchers 3 and 4 highlights the influence of research background on the results of downstream analyses. Researchers 1, 2 and 5 showed no statistical significance from Livecyte segmented cells in terms of cell area, but were statistically significant for both circularity and mean gray value. In contrast, researchers 3 and 4 showed no statistical significance from Livecyte segmented cells in terms of circularity, but were statistically significant for both area and mean gray value.

The impact of inter- and intra-user variability in extracted metrics was further investigated using PCA. Scores plots for cells 1, 2 and 3 are provided in **Figure 3(b) i, ii** and **iii** respectively. Researcher-specific clusters can be identified within the scores plots demonstrating idiosyncrasies in cell segmentation and their impact on the resulting extracted metrics. Clusters from researchers 1, 2 and 5 tended to overlap with one another as did the clusters from researchers 3 and 4, again suggesting a relationship between research experience and the results obtained. Intra-user variability is visible within all three scores plots, with larger ellipses representative of greater variability. Tighter clusters were frequently obtained with Livecyte, even in cases where all researchers displayed extensive variability. This is the case for cell 3, where greater consistency in results is obtained with Livecyte across all three cells.

Additionally, intra-user variability in extracted metrics was quantified through the calculation of Euclidean distances to measure the pair-wise similarity of data vectors obtained by each researcher, the results are shown in **Figure 3(c)**. ANOVA identified statistical significant differences as a result of cell number (*p* value = 0.0273), indicating that certain cells invoked greater variability in extracted metrics in comparison to others. A statistically significant difference was also identified between researchers, demonstrating that some were more consistent in their segmentation approach and this was reflected in the consistency of the extracted metrics. The interaction between cell number and researcher was also highly statistically significant, indicating that consistency posed a greater challenge for some cells for certain researchers, resulting in greater variability in extracted metrics for some cells. Notably the distances calculated for Livecyte segmentation remained consistent across all three cells, with the distances themselves minimised in comparison to those obtained by other researchers indicating greater consistency in extracted metrics as a result of Livecyte segmentation.

### Inter-user variability in cell tracking

As well as segmenting each of the three cells 5 times, the researchers tracked cell 1, 2 and 3 for a total of 50 frames three times each to assess inter-user variability in cell tracking. Note that the researcher previously referred to as researcher 3 did not take part in this cell tracking study and was replaced with a different researcher referred to as researcher 6, with similar expertise to researcher 3. Researchers selected the cell’s position on consecutive frames to form three, 50-frame long trajectories. The inter-user variability in cell trajectories can be observed in **Figure 4(a)**, where, for each cell, the researchers’ first tracking attempt is shown. By eye, the general shape of trajectories obtained for cells 1 and 2 appear fairly consistent, with subtle differences along the trajectories induced by variability in click position of each researcher. The trajectory shape for cell 3, shows a noticeable difference for researchers 4 and 6 in comparison to researchers 1, 2 and 5 although the trajectories produced by researchers 4 and 6 are similar.

**Figure 4:**
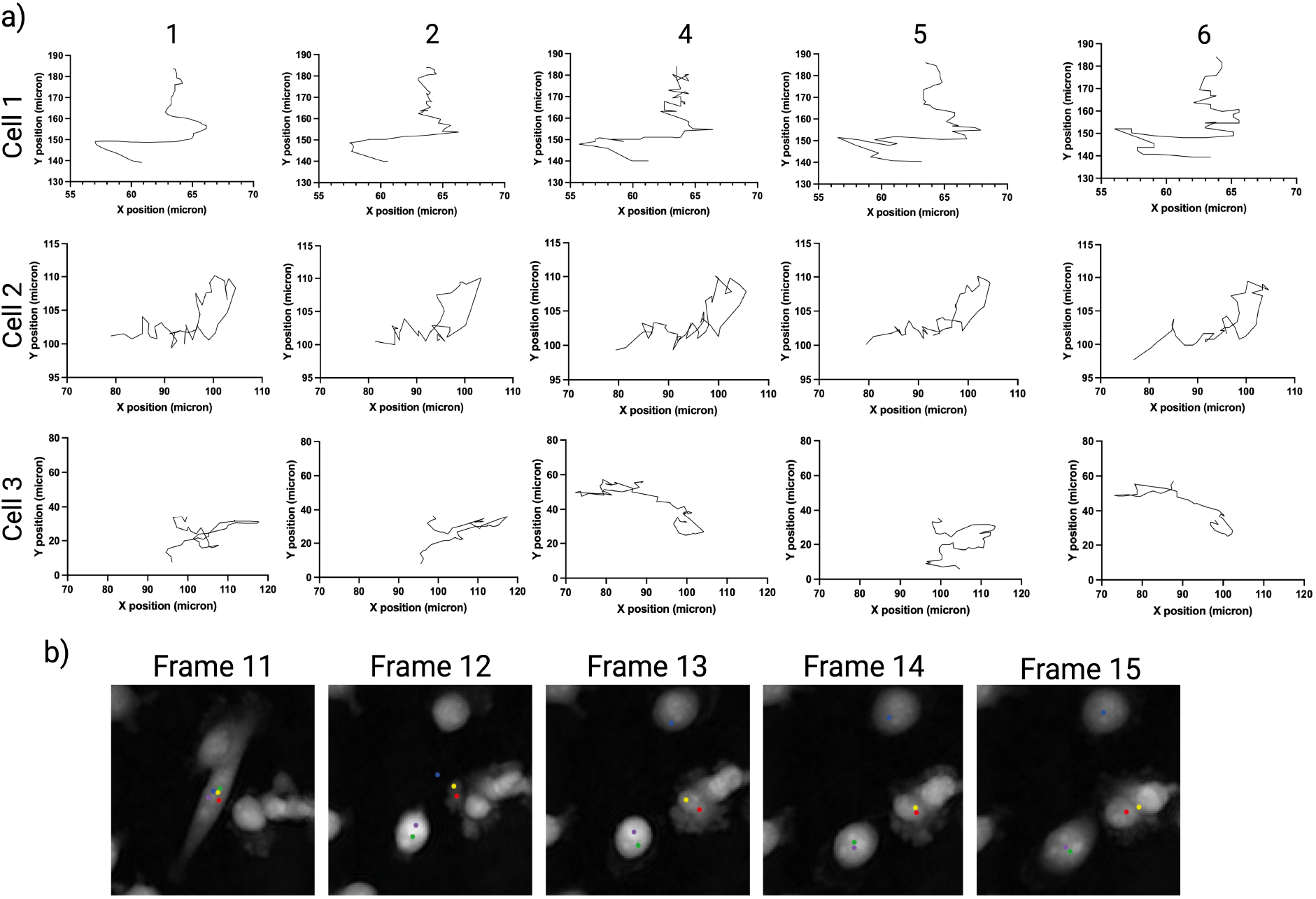
Inter-user variability in cell tracking. **(a)** Trajectories from each researcher’s first cell tracking of cells 1, 2 and 3. For cells 1 and 2, subtle differences between trajectories can be observed, induced by variability in click position between researchers. More notable inter-user variability can be observed for cell 3, where differences in cell identification between researchers resulted in the tracking of different cells that were misidentified as cell 3. **(b)** Visualisation of the misidentification of cell 3 throughout the time-lapse. All researchers correctly identify cell 3 in frame 11, where points represent click position coloured by researcher. Cell 3 undergoes a change in morphology on frame 12 which causes disagreement on which cell is actually cell 3. Researchers 4 and 6 correctly identify cell 3 on frame 12, researchers 2 and 5 misidentify a different cell as cell 3 and researcher 1 identifies background as cell 3. Each researcher then continues to track their chosen cells for the remainder of the time-lapse, with researcher 1 misidentifying a cell as cell 3 in frame 13.

Further investigation showed that the difference is due to disagreement in cell identification from one frame to the next, as visualised in **Figure 4(b)**. All researchers correctly identify cell 3 in frame 11, as demonstrated by all 5 points within the cell interior. The cell then undergoes a notable change in morphology on frame 12, making it correct identification of cell 3 challenging on this frame. Cell 3 is correctly identified by researchers 4 and 6 (green and purple points) in frame 12 whereas researchers 2 and 5 (red and yellow points) misidentify a different cell as cell 3. Researcher 1 (blue point) selects a position that is actually background as opposed to a cell interior on frame 12. This could be due to an unexpected change in morphology, and therefore position, of cell 3 between frames 11 and 12. Researcher 1 does select a cell interior on frame 13 but again this is a misidentification rather than cell 3. All researchers then continue to track their selected cells for the remainder of the time-lapse, resulting in different trajectories with only researchers 4 and 6 correctly tracking the original cell 3.

### Intra-user variability in cell tracking

As the main source of variation in cell tracking arises from inconsistencies in researcher click positions, we sought to explore the intra-user variability present across three independent cell tracking attempts for each cell. We quantified each researcher’s click spread for each of the three cells and scatter plots displaying these values are provided in **Figure 5(a)**. Here the origins are representative of the average click position from three tracking attempts, with the click spread of each point then calculated as its deviation from the average. Researchers who remain consistent in their click positions will have all points closely clustered and centred around the origin. Researchers with inconsistencies in click position will have deviated scatter plots with points positioned at varying distances from the origin. All researchers experienced intra-user variability in click position across all cells, with researcher 6 providing the largest inconsistencies for cells 1 and 2 as evidenced by greater deviation in personal click spread for these cells.

**Figure 5:**
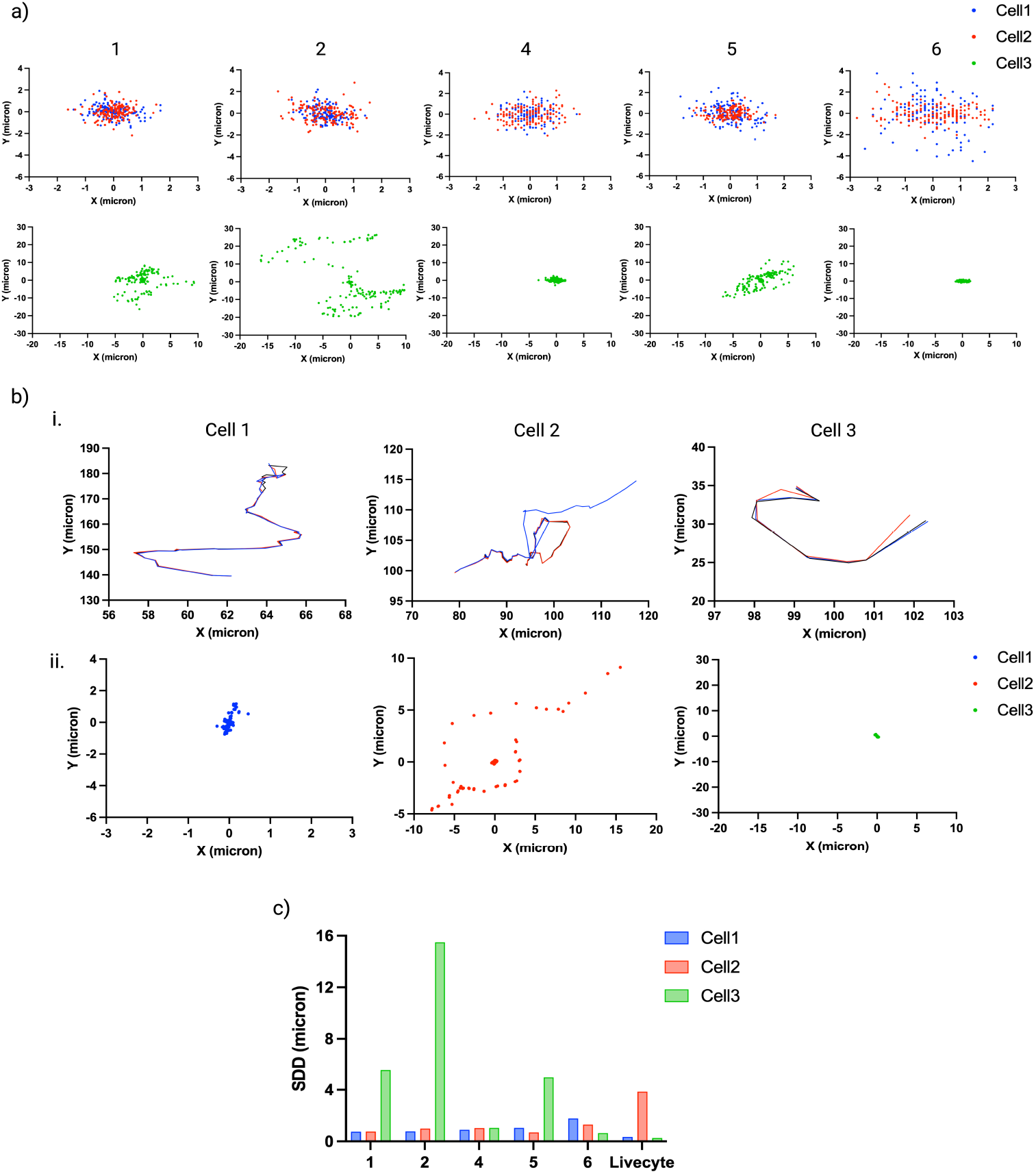
Intra-user variability in cell tracking. **(a)** Click spread plots for each researcher, coloured by cell number. Origins represent the average click position across three tracking attempts, with the click spread of each point calculated as deviation from this average. Click spread plots for cell 3 show greater deviance due to repeated misidentification of cell 3 throughout the time-lapse resulting in variable cell trajectories. **(b)** i. Cell trajectories for cell 1, 2 and 3 obtained from automated tracking using Phasefocus CATbox. Each colour represents one of three trajectories obtained from the same cell time-lapse independently processed to replicate inter-user variability. Misidentification of cell 2 in one time-lapse results in an inaccurate trajectory (shown in blue). Trajectories for cell 3 are shorter than those obtained by manual tracking due to the premature ending of tracking on frame 11 when cell 3 undergoes a morphology change, after which the true cell 3 is given a new cell identifier for the remainder of the time-lapse. ii. Centroid spread plots for each researcher, coloured by cell number. Origins are represent the average centroid position across each of three tracking repeats, with the spread of each centroid calculated as deviation from this average. Deviation is minimal for cell 1 and cell 3, but the misidentification of cell 2 in one trajectory causes deviation in centroids. **(c)** Standard distance deviation (SDD) values to quantify the manual click spread from **(a)** and automated centroid spread from **(b)**.

Click spread for cell 3 was considered separately to cells 1 and 2 due to the impact of frequent misidentification of cell 3. Researchers 4 and 6, the two researchers who successfully identified cell 3 in **Figure 4(b)** successfully identified cell 3 in all three of their tracking attempts, resulting lower click spread for cell 3 in comparison to all other researchers. All other researchers identified different cells as cell 3 within their three tracking attempts, resulting in large click spreads for researchers 1, 2 and 5 for cell 3. Interestingly these results suggest a relationship between cell tracking results and research expertise, with cell biologists 4 and 6 displaying greater accuracy in cell tracking in comparison to microscopists 1, 2 and 5.

In the cases of cell 1 and 2, the click spread deviation for each researcher is likely a result of differences in determination of the cell centroid. Researchers with low click spread potentially aim for the same sub-cellular region on each attempt, whereas high click spread indicates inconsistency in selection of sub-cellular region. The click spreads for cell 3 are dominated by the challenge of identifying cell 3 when it experiences a change in morphology on frame 12 with the additional deviation as a result of variability in selection of a sub-cellular region.

We next sought to compare manual tracking results with those obtained from automated tracking produced by the Phasefocus CATbox. Replicates were obtained by tracking the cells after three independent image processing pipelines were performed. Different processing methods were found to affect the determination of cell centroids. The obtained cell trajectories, together with centroid deviation, for each cell are provided in **Figure 5(b)**. The plots show that despite different image processing pipelines used, the tracking results are consistent for cell 1 with negligible variation in the cell trajectories and very little deviation in centroid position with points in the scatter plot forming a tight cluster centered at the origin. The same can be seen for cell 3 although each track ends at frame 11, just before the cell changes morphology on frame 12 where each tracking repeat generated a new identifier for the cell that was used for the remainder of the time-lapse. There was notable misidentification of cell 2 in one tracking repeat as evidenced by a change in the cell trajectory and centroid deviation, shown for a subset of points within the centroid scatter plot.

Standard distance deviation (SDD) was used to quantify the overall deviation of points for each cell and each researcher (**Figure 5(c)**). This confirmed observations from the click spread plots with researcher 6 displaying greater inconsistencies in click positions for cell 1 and 2 in comparison to all other researchers. Variability in SDD was greatest for cell 3 due to repeated misidentification of cell 3 by multiple researchers. SDD was minimal for Livecyte tracking of cells 1 and 3, but higher than all other researchers for cell 2. These results highlight greater consistency and reproducibility of automated tracking in comparison to manual tracking but also highlight problems in automated cell tracking.

### The impact of cell tracking on extracted metrics

As inter- and intra-user variability in manual segmentation was found to have a major impact on morphological metrics, we hypothesised that the same would be the case for extracted characteristic metrics of cell trajectories. Variability in cell trajectories, both between researchers and within each researcher’s own repeats, occurs due to variability in click positions on consecutive frames. Certain dynamical features, such as speed of cells, are extremely sensitive to such deviations resulting in large variability in the time series obtained for these features (**Figure 6(a)i, ii**). The variability across the whole cell trajectory has a cumulative effect on summary statistics that are calculated to characterise cell behaviour throughout the whole time-lapse (such as mean speed), with each researcher obtaining different values dependent on their own tracking approach. The mean speeds calculated from each researchers’ cell 1 trajectories are shown in **Figure 6(a)iii**. ANOVA revealed statistically significant differences in mean speed between researchers performing the tracking (*p* value = 0.0017), highlighting the impact of subjectivity on quantification. Notably, the minimal standard error of the mean was achieved by automated Livecyte tracking (s.e = 0.034) in comparison to manual tracking (s.e = 0.057, 0.054, 0.088, 0.12 for researchers 1, 2, 4 and 6 respectively). PCA was used to assess the cumulative effect of such variability on a combination of motility metrics. As for PCA of morphological metrics shown in **Figure 3(b)**, researcher-specific clusters can be identified within the scores plots demonstrating idiosyncrasies in cell tracking and the impact these have on the resulting extracted metrics. Intra-user variability is visible within all three scores plots, with larger ellipses representative of greater variability. Largest ellipses within scores plots for cell 1 and 2 are observed for researcher 6, found to have the largest click spread deviation for these cells. Automated Livecyte tracking resulted in the smallest ellipse, and therefore the least variability in motility metrics, for cell 1 but the impact of misidentification of cell 2 can be observed in the cell 2 scores plot where a much larger Livecyte ellipse and more dispersed points can be observed. Note that Livecyte points are not included in the scores plot for cell 3 due to the premature ending of all 3 Livecyte trajectories for this cell meaning metrics such as track length could not be compared with full length manual trajectories.

**Figure 6:**
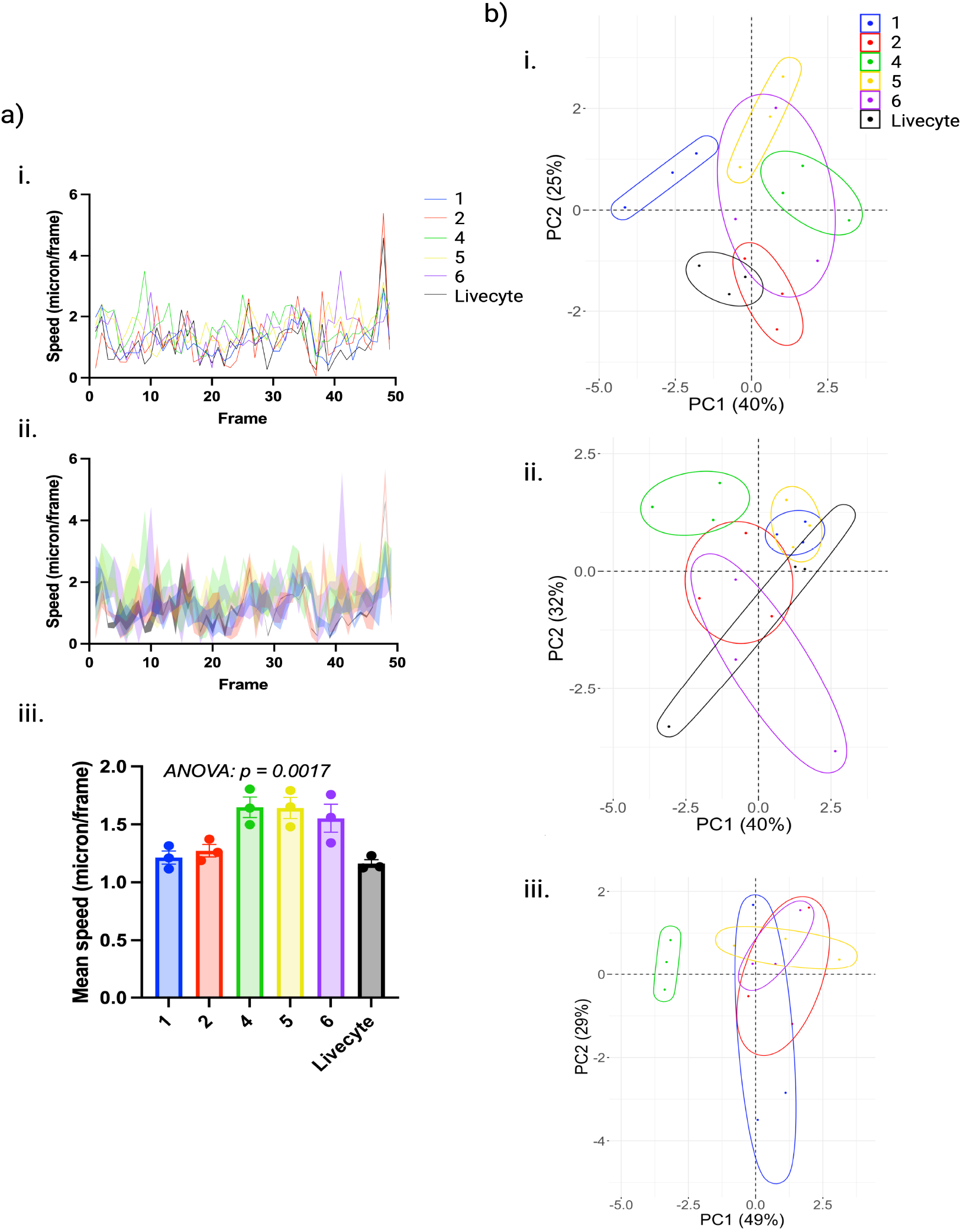
The impact of cell tracking on extracted metrics. **(a)** i. Time series for mean speed obtained from three tracking attempts for cell 1, coloured by researcher. ii. The same plot as in i but with the region between SEM error bars shaded to display intra-user inconsistencies. iii. Separate scatter plots showing mean speeds calculated for each researcher and each tracking repeat of cell 1. ANOVA revealed a statistically significant relationship between mean speed and the researcher performing tracking (*p* value = 0.0017). **(b)** The resulting PCA scores plots for i. cell 1, ii. cell 2 and iii. cell 3, performed with motility metrics extracted from the trajectories. Points are coloured according to researcher and 95% confidence ellipses are displayed. Small and large ellipses are representative of low and high intra-user variability in extracted metrics respectively. Note that Livecyte points are not included in the scores plot for cell 3 due to the premature ending of all 3 Livecyte trajectories.

### Variability in extracted metrics from segmentation software

Cell segmentation software were benchmarked on their ability to obtain accurate cell ROIs that reliably capture the true morphology of the cells within the images. For this analysis, a cell image was manually segmented by an expert with extensive experience working with ptychographic images as well as the MDA-MB-231 cells within the image (**Figure 7a**). Manually segmented cells were used as a gold standard, with the accuracy of automated software quantified by how closely the outputs matched those from manual segmentation. From the automated segmentation results presented within **Supplementary Figure 1** it can be observed that morphology of ROIs from FIJI and Icy, in particular, differ from those of the other 3 software. Notably, FIJI and Icy tend to segment the cell body, excluding the lower intensity cell protrusions that are identified by Livecyte, CellProfiler, CellPose and manual segmentation. PCA of all morphological metrics extracted from each software’s obtained ROIs confirmed this to be true, with FIJI and Icy ellipses providing smallest overlap with the manual segmentation ellipse and greatest distance between group means in the PCA scores plot (**Figure 7b**).

**Figure 7:**
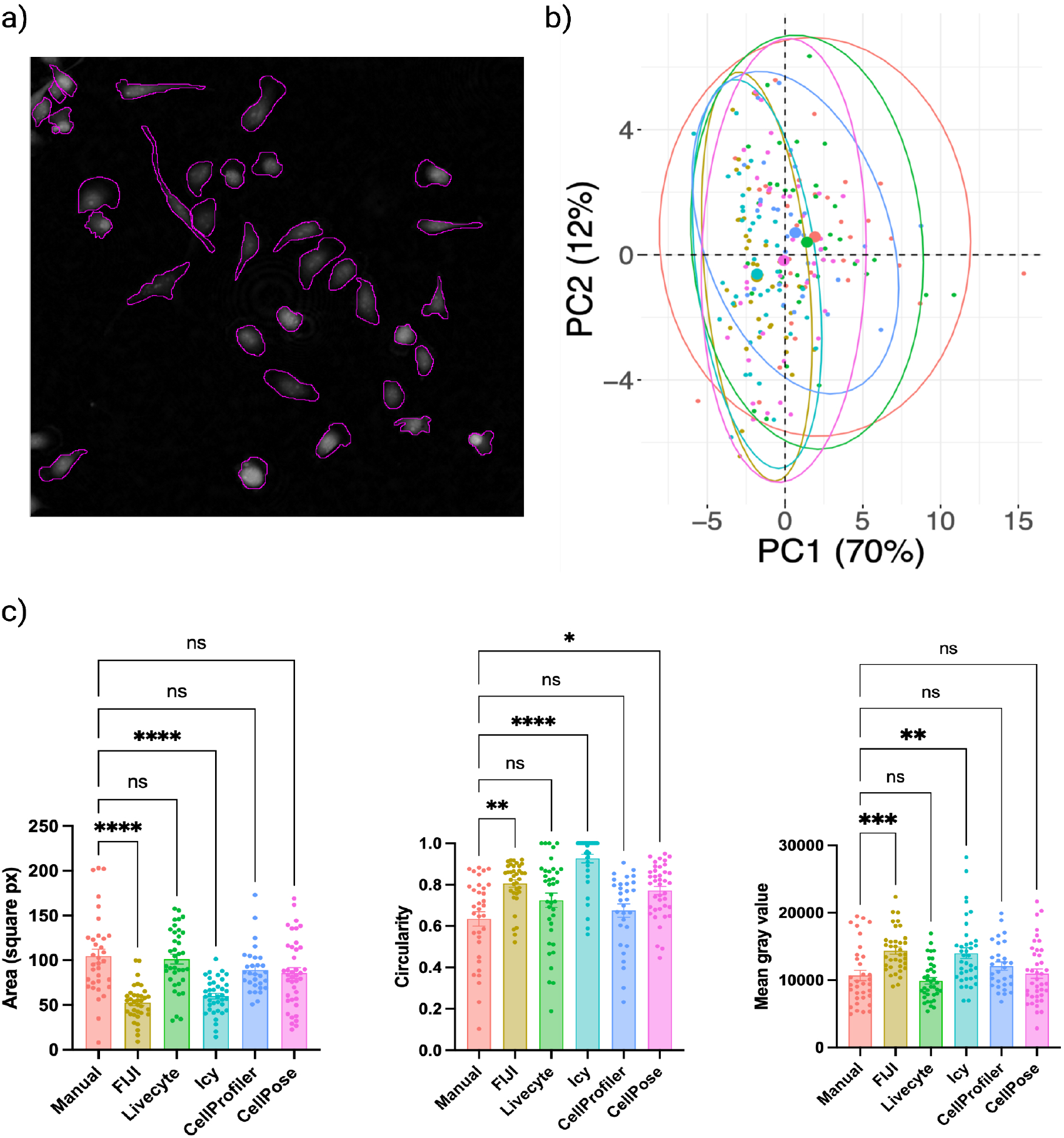
Variability in extracted metrics from segmentation software. **(a)** Results of manual segmentation on the 0h cell image, performed by an expert with extensive experience working with ptychographic images and the MDA-MB-231 cell line used in this study. **(b)** PCA of morphological metrics extracted from cell ROIs, coloured by segmentation approach. FIJI and Icy display greatest variability to manual segmentation with their ellipses sharing the minimum overlap with the manual segmentation ellipse. Larger points are representative of group means. **(c)** Distributions for cell area, circularity and mean gray value extracted by manual segmentation and five different automated software packages. Bars show group means and error bars represent SEM. Results of Dunn’s multiple comparisons test are displayed on plots, with greatest significance across all metrics obtained by FIJI and Icy segmented cells when compared with manual segmentation.

Further investigation into the morphological metrics impacted by this difference in segmentation showed that FIJI and Icy segmentation result in cells that are significantly smaller in size, have increased circularity and have higher mean gray value compared to manually segmented cells (**Figure 7c**). In comparison, Livecyte, Cell-Profiler and CellPose segmentation resulted in the extraction of morphological metrics that were not significantly different to those obtained by manual segmentation, the exception being circularity of CellPose ROIs (*p* value = 0.046).

## Discussion

The results presented here demonstrate the importance of choosing a segmentation and tracking approach that is optimised for the application in question. Most segmentation and tracking software rely on the contrast in intensity between cells and background to accurately detect cell boundaries. These software are often optimised for fluorescence images where fluorescent labels enhance the intensity of pixels within cell interiors. With pytchography being a relatively new imaging modality, pre-existing algorithms do not perform as well on these images due to differences in the distribution of cell pixel intensities within label-free images. The newer segmentation and tracking software, Livecyte and CellPose-TrackMate, were found to be optimal for our images in comparison to the other, well-established software tested: FIJI, Icy and CellProfiler. Livecyte algorithms are optimised for ptychographic imaging and this shows in its superiority for both segmentation and tracking. CellPose is the most recent software within the field and already offers several pre-built models that are optimised for imaging modalities, namely phase-contrast and brightfield, that often pose the greatest challenge for segmentation.^16^ As label-free imaging becomes more established, it is likely that pre-existing segmentation and tracking software will further develop their current algorithms to handle such images in order to compete with newer approaches.

Our findings suggest that the results of manual segmentation and tracking should be used as a gold standard with caution. Deep learning algorithms frequently make use of manual results for training and testing performance, yet manual results vary significantly with experience and subjectivity. It is important that manual segmentation and tracking for such ground truth data sets is performed by an expert in the particular field in question, and that their own results provide minimal intra-user variability. The subjectivity of manual segmentation is of particular concern for medical applications in which segmentation results are used for diagnosis of diseases, such as the detection of breast tumours in mammograms. Furthermore, the morphological metrics extracted from segmented regions are used to assess response to treatment and disease progression (such as size of tumour following chemotherapy). This motivates the need for standardised, automated approaches to segmentation that eradicate the potential for bias within results.

The variability in segmentation and tracking results, both inter- and intra-user as well as inter-software support the ongoing replicability and reproducibility crisis within scientific studies. The results of this study showed that idiosyncrasies present within initial segmentation and tracking are further exacerbated by the extraction of metrics to quantify cell morphology and motility. This led to results from identical cell populations and even same cells being found statistically significantly different based on their initial segmentation and tracking. Automated software takes on an algorithmic approach to cell segmentation and tracking, following the same formulaic pipeline both within and between images. On the other hand, researchers performing manual segmentation and tracking often do not have a set of rules to follow in order to detect cell boundaries or centroids, as evidenced by intra-user variability in identified ROIs and researcher click spread, therefore adding cumulative inaccuracies to their results. Intra-software variability was minimal which motivates the use of automated approaches to improve the repeatability of results. However, though results may be more reproducible this does not ensure that they are accurate. It is therefore vital that background research is done on the choice of automated segmentation and tracking software prior to downstream analysis.

The similarity of results from manual segmentation and tracking tend to be associated with the similarity of research background. Notably, the trained microscopists (researchers 1, 2 and 5) produced manual segmentation results with greater similarity to each other than to cell biologists (researchers 3 and 4). This was supported by the calculation of Dice coefficients for quantification of segmentation overlap. This suggests that research background is a latent variable in terms of quantification from cell segmentation and tracking, though this impact is often not assessed or accounted for in scientific research.

The challenge caused by cell 3, even for trained microscopists and automated software, emphasises the importance of minimising interval times between frames during image acquisition. Cells that undergo a drastic morphology change from one frame to another are difficult to identify in consecutive frames, and misidentification results in erroneous changes to trajectories and consequently the time series of extracted metrics. These results demonstrate that both misidentification and erroneous cell segmentation are visible within extracted morphology and motility metrics and suggest the possibility of automating detection of such instances to clean up data sets prior to downstream analyses. Shorter cell tracks would not cause a problem in terms of biological conclusions, as the remaining cell track is still represented within the data set but linked to a different cell identifier. To handle the longer, inaccurate cell tracks, machine learning models could be used to identify interruptions in cell time series induced by misidentification and exclude these cells from further analysis, a step that is included in the CellPhe toolkit for automated cell phenotyping.^26^

## Acknowledgements

This work was supported by the BBSRC (grant number B/S507416/1). The authors thank the following members of the research team for assisting with data collection: Dr. Serife Yerlikaya, Nattanan Sajjaboontawee, Jodie Malcolm, and Blythe Wright from the University of York, and Kundi Umar and Abdurrahman Alkham from Yobe State University.

## References

1 W. Chen, W. Li, X. Dong, et al. A review of biological image analysis. Current Bioinformatics, 12, 2017.

2 Y. Al-Kofahi, A. Zaltsman, R. Graves, et al. A deep learning-based algorithm for 2-d cell segmentation in microscopy images. BMC bioinformatics, 19(1):1–11, 2018.

3 Y. Zhang, P. Zhong, D. Jie, et al. Brain tumor segmentation from multi-modal mr images via ensembling unets. Frontiers in Radiology, page 11, 2021.

4 P. Bándi, R. Van de Loo, M. Intezar, et al. Comparison of different methods for tissue segmentation in histopathological whole-slide images. In 2017 IEEE 14th International Symposium on Biomedical Imaging (ISBI 2017), pages 591–595. IEEE, 2017.

5 F. Dionisio, L. S. Oliveira, M. Hernandes, et al. Manual versus semiautomatic segmentation of soft-tissue sarcomas on magnetic resonance imaging: evaluation of similarity and comparison of segmentation times. Radiologia Brasileira, 54:155–164, 2021.

6 E. Covert, K. Fitzpatrick, J. Mikell, et al. Intra-and inter-operator variability in mri-based manual segmentation of hcc lesions and its impact on dosimetry. EJNMMI physics, 9(1):1–16, 2022.

7 L. Joskowicz, D. Cohen, N. Caplan, et al. Inter-observer variability of manual contour delineation of structures in ct. European radiology, 29:1391–1399, 2019.

8 H. K. Bø, O. Solheim, A. S. Jakola, K. Kvistad, et al. Intra-rater variability in lowgrade glioma segmentation. Journal of Neuro-oncology, 131:393–402, 2017.

9 N. Sharma, L. Aggarwal, et al. Automated medical image segmentation techniques. Journal of medical physics, 35(1):3, 2010.

10 H. Chan, L. M. Hadjiiski, and R. K. Samala. Computer-aided diagnosis in the era of deep learning. Medical physics, 47(5):e218–e227, 2020.

11 R. A. Morey, C. M. Petty, Y. Xu, et al. A comparison of automated segmentation and manual tracing for quantifying hippocampal and amygdala volumes. Neuroimage, 45(3):855–866, 2009.

12 K. Verma, S. Kumar, and D. Paydarfar. Automatic segmentation and quantitative assessment of stroke lesions on mr images. Diagnostics, 12(9):2055, 2022.

13 J. Walsh, A. Othmani, M. Jain, et al. Using u-net network for efficient brain tumor segmentation in mri images. Healthcare Analytics, 2:100098, 2022.

14 V. Wiesmann, D. Franz, C. Held, et al. Review of free software tools for image analysis of fluorescence cell micrographs. Journal of Microscopy, 251:39–53, 2015.

15 C. Edlund, T. R. Jackson, N. Khalid, et al. Livecell—a large-scale dataset for label-free live cell segmentation. Nature methods, 18(9):1038–1045, 2021.

16 C. Stringer and M. Pachitariu. Cellpose 2.0: how to train your own model. BioRxiv, pages 2022–04, 2022.

17 J. R. Masters et al. Short tandem repeat profiling provides an international reference standard for human cell lines. Proceedings of the National Academy of Sciences, 98(14):8012–8017, 2001.

18 M. Yang, D. J. Kozminski, L. A. Wold, et al. Therapeutic potential for phenytoin: targeting nav1.5 sodium channels to reduce migration and invasion in matastatic breast cancer. Breast Cancer Research and Treatment, 134(2):603–615, 2012.

19 D. J. McLean, Skowron V., and A. Marta. trajr: An r package for characterisation of animal trajectories. Ethology, 124(6):440–448, 2018.

20 J. Schindelin, I. Arganda-Carreras, E. Frise, et al. Fiji: an open-source platform for biological-image analysis. Nature Methods, 9:676–682, 2012.

21 Punam K. Saha, Felix W. Wehrli, and Bryon R. Gomberg. Fuzzy distance transform: Theory, algorithms, and applications. Computer Vision and Image Understanding, 86:171–190, 2002.

22 F. De Chaumont, S. Dallongeville, N. Chenouard, et al. Icy: an open bioimage informatics platform for extended reproducible research. Nature methods, 9:690–696, 2012.

23 C. McQuin, A. Goodman, V. Chernyshev, et al. Cellprofiler 3.0: Next-generation image processing for biology. 16:2018.

24 C. H. Li and C. K. Lee. Minimum cross entropy thresholding. Pattern Recognition, 26:617–625, 1993.

25 C. Stringer, T. Wang, M. Michaelos, et al. Cellpose: a generalist algorithm for cellular segmentation. Nature methods, 18(1):100–106, 2021.

26 L. Wiggins, A. Lord, K. Murphy, et al. The cellphe toolkit for cell phenotyping using time-lapse imaging and pattern recognition. Nature Communications, 14(1):1854, 2023.

