## Supplementary Data for "Exploring the impact of variability in cell segmentation and tracking approaches"

**Table S1: Results of Tukey's multiple comparisons test for area**

| **Comparison** | **Summary** | $p$ **value** |
|---|---|---|
| User1 vs. User2 | ns | 0.9762 |
| User1 vs. User3 | **** | <0.0001 |
| User1 vs. User4 | **** | <0.0001 |
| User1 vs. User5 | ns | >0.9999 |
| User1 vs. Livecyte | ns | 0.1258 |
| User2 vs. User3 | **** | <0.0001 |
| User2 vs. User4 | **** | <0.0001 |
| User2 vs. User5 | ns | 0.9338 |
| User2 vs. Livecyte | ns | 0.4745 |
| User3 vs. User4 | ns | 0.8724 |
| User3 vs. User5 | **** | <0.0001 |
| User3 vs. Livecyte | **** | 0.3630 |
| User4 vs. User5 | **** | <0.0001 |
| User4 vs. Livecyte | **** | 0.6090 |
| User5 vs. Livecyte | ns | 0.0785 |

**Table S2: Results of Tukey's multiple comparisons test for circularity**

| **Comparison** | **Summary** | $p$ **value** |
|---|---|---|
| User1 vs. User2 | ns | 0.0904 |
| User1 vs. User3 | **** | <0.0001 |
| User1 vs. User4 | **** | <0.0001 |
| User1 vs. User5 | ns | 0.9991 |
| User1 vs. Livecyte | **** | <0.0001 |
| User2 vs. User3 | **** | <0.0001 |
| User2 vs. User4 | **** | <0.0001 |
| User2 vs. User5 | * | 0.0356 |
| User2 vs. Livecyte | **** | <0.0001 |
| User3 vs. User4 | ns | 0.9988 |
| User3 vs. User5 | **** | <0.0001 |
| User3 vs. Livecyte | ns | 0.3630 |
| User4 vs. User5 | **** | <0.0001 |
| User4 vs. Livecyte | ns | 0.6090 |
| User5 vs. Livecyte | **** | <0.0001 |

**Table S3: Results of Tukey's multiple comparisons test for mean gray value**

| **Comparison** | **Summary** | $p$ **value** |
|---|---|---|
| User1 vs. User2 | ns | 0.1615 |
| User1 vs. User3 | **** | <0.0001 |
| User1 vs. User4 | **** | <0.0001 |
| User1 vs. User5 | ns | 0.9975 |
| User1 vs. Livecyte | **** | <0.0001 |
| User2 vs. User3 | **** | <0.0001 |
| User2 vs. User4 | **** | <0.0001 |
| User2 vs. User5 | ns | 0.0562 |
| User2 vs. Livecyte | * | 0.0109 |
| User3 vs. User4 | ns | >0.9999 |
| User3 vs. User5 | **** | <0.0001 |
| User3 vs. Livecyte | **** | <0.0001 |
| User4 vs. User5 | **** | <0.0001 |
| User4 vs. Livecyte | **** | <0.0001 |
| User5 vs. Livecyte | **** | <0.0001 |

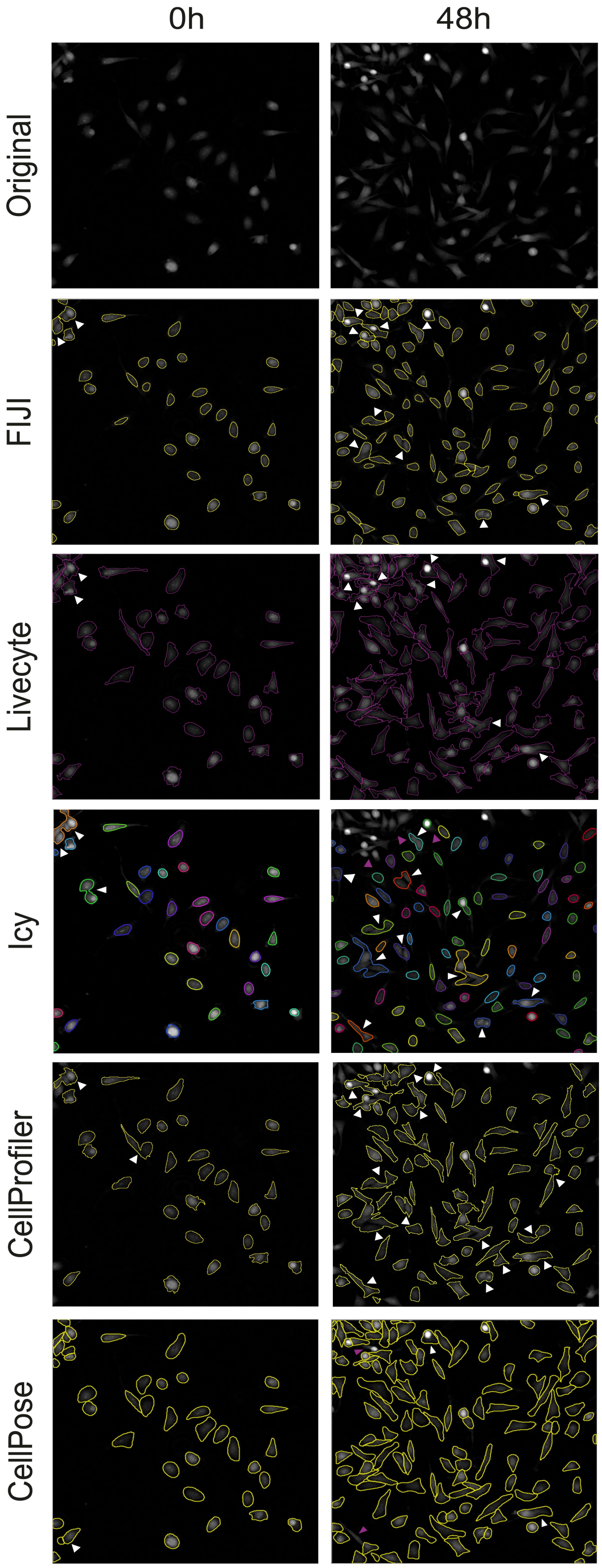




**Figure S1: Segmentation results from a panel of automated segmentation software**

Automated segmentation results obtained from FIJI, Livecyte, Icy, CellProfiler and CellPose on two cell images taken at 0 hours and 48 hours of a time-lapse. Instances of over- and undersegmentation are highlighted by white arrows, and cases where a cell is unidentified by automated segmentation are highlighted by pink arrows.
